# Route-dependent spatial engram tagging in mouse dentate gyrus

**DOI:** 10.1101/2022.06.20.496824

**Authors:** Lucius K. Wilmerding, Ivan Kondratyev, Steve Ramirez, Michael E. Hasselmo

## Abstract

The dentate gyrus (DG) of hippocampus is hypothesized to act as a pattern separator that distinguishes between similar input patterns during memory formation and retrieval. Sparse ensembles of DG cells associated with learning and memory, i.e. engrams, have been labeled and manipulated to recall novel context memories. Functional studies of DG cell activity have demonstrated the spatial specificity and stability of DG cells during navigation. To reconcile how the DG contributes to separating global context as well as individual navigational routes, we trained mice to perform a delayed-non-match-to-position (DNMP) T-maze task and labeled DG neurons during performance of this task on a novel T-maze. The following day, mice navigated a second environment: the same T-maze, the same T-maze with one route permanently blocked but still visible, or a novel open field. We found that the degree of engram reactivation across days differed based on the traversal of maze routes, such that mice traversing only one arm had higher ensemble overlap than chance but less overlap than mice running the full two-route task. Mice experiencing the open field had similar ensemble sizes to the other groups but only chance-level ensemble reactivation. Ensemble overlap differences could not be explained by behavioral variability across groups, nor did behavioral metrics correlate to degree of ensemble reactivation. Together, these results support the hypothesis that DG contributes to spatial navigation memory and that partially non-overlapping ensembles encode different routes within the context of different environments.

**Highlights:**

- Immediate-early-gene labeling strategy revealed spatial navigation ensembles in DG
- Sub-ensembles encode separate maze routes within a larger task context
- Ensemble reactivation does not correlate with behavioral variables

## 1. INTRODUCTION

The dentate gyrus (DG), a subregion of the hippocampal formation, is hypothesized to act as a pattern separator that distinguishes between similar input patterns during memory formation and retrieval (Marr, 1971; McNaughton & Morris, 1987; O’Reilly & McClelland, 1994; Treves & Rolls, 1994; Hasselmo & Wyble, 1997; Leutgeb et al., 2007; Neunuebel & Knierim, 2014). Sparse ensembles of DG memory-associated granule cells, or engram cells, have been optogenetically targeted to successfully influence memory-associated behavior (Liu et al., 2012; Ramirez et al., 2013; Denny et al., 2014; Redondo et al., 2014) even after consolidation (Kitamura et al., 2017) and pathogenic aging leading to natural recall failure (Roy et al., 2016). These findings support the hypothesis that the DG encodes the contextual dimension of memories. Another body of literature emphasizes the spatial representations of DG granule cells, which exhibit place fields similar to those in CA1 (O’Keefe & Dostrovsky, 1971; Neunuebel & Knierim, 2012, 2014; GoodSmith et al., 2017; Hainmueller & Bartos, 2018; Cholvin et al., 2021). Lesions of the DG granule cell population impair spatial memory (McLamb et al., 1988; McNaughton et al., 1989; Nanry et al., 1989; Xavier et al., 1999; for a review, see Xavier & Costa, 2009) and reduce both spatial specificity of CA3 place cells and reward-associated sharp-wave-ripple rate (Sasaki et al., 2018). Reconciling how the DG contributes to both novel contextual learning and spatial representations would offer important insight into the contribution of this region for learning and memory.

To that end, our study investigated the role of the DG in distinguishing between multiple routes within a single T-maze context during a spatial navigation task. Male and female mice were trained on a delayed-non-match-to-position (DNMP) task with two routes. A population of active DG granule cells was visualized using an immediate-early-gene strategy of labeling cFos positive cells active during exposure on Day 1 to a novel 2-route T-maze (Guzowski et al., 1999; Reijmers et al., 2007; Liu et al., 2012; Ramirez et al., 2013). Another population of cFos positive cells activated on Day 2 by a second behavioral context was visualized with immunohistochemical staining for comparison. We hypothesized that the DG of mice exposed to a 1-route maze on Day 2 would show more overlap between the two cell populations than chance, but less overlap than mice re-exposed to the full 2-route maze from Day 1. Our findings support this hypothesis and additionally reveal the size and degree of ensemble reactivation are largely independent of behavioral performance in the arenas. Our results indicate that the DG plays a role in encoding particular sub-routes of a 2D environment during ongoing spatial navigation.

## 2. METHODS

### 2.1 Subjects

25 wildtype (WT) C57B6J male and female mice (Jax) were segregated by sex and group-housed prior to surgery. They received food and water ad libitum and were placed on a diet containing 40 mg/kg doxycycline (dox; Bio-Serv) at least 1 week prior to surgery at the age of 17-36 weeks. Post-surgically, the mice were housed in pairs (by sex) on a reverse 12 hour light-dark cycle. Mice were split equally into groups using random assignment, counterbalancing for sex. All procedures related to mouse care and treatment were in accordance with Boston University and National Institutes of Health guidelines for the Care and Use of Laboratory animals.

### 2.2 Viral constructs and packaging

The AAV-cFos-tTA and AAV-cFos-tTa-TRE-eYFP were constructed as described previously (Ramirez et al., 2013) and sourced from Gene Therapy Center and Vector Core at the University of Massachusetts Medical School. The viral titrations were 1.5*10^13^ genome copy per mL for AAV-cFos-tTA-TRE-eYFP and 1.5*10^13^ genome copy per mL for AAV-cFos-tTA.

### 2.3 Stereotactic injection

All surgeries were performed under stereotactic guidance and all coordinates are reported relative to bregma. Anesthesia was induced with 5.0% isoflurane and maintained thereafter at a concentration of 1.5-2.0%. Bilateral burr holes were made at -2.2 mm (AP) and +/-1.3 (ML) using a 0.5mm drill to allow a 10uL nanofil syringe (World Precision Instruments) to be lowered to 2.0 mm (DV). 300nL of AAV virus cocktail was injected bilaterally at a rate of 100nL/min controlled by a MicroSyringe Pump Controller (World Precision Instruments). Following injection, the needle was kept at the injection site for 5 minutes and slowly withdrawn afterwards. Bone wax was gently inserted into the burr holes to seal the skull, and two to three sterile sutures were used to close the wound. Post-operative subcutaneous injections of Buprenorphine (0.1 mg/kg) and Ketoprofen (5 mg/kg) were administered for three days following surgery and Enrofloxacin (10 mg/kg) for five days post-surgically.

### 2.4 Behavioral assays and engram tagging

All behavioral assays were performed during the light cycle of the day (7:00 - 18:00) on animals 17-32 weeks old. Mice were handled 2-5 min per day for two days before behavioral training. To increase training motivation, mice were water restricted during the training period, tagging and re-exposure period to 20 minutes of *ad libitum* water access daily, in addition to sucrose consumed in the maze.

Behavior was run in a two-arm T-maze, as previously described (Fig. 1; Levy et al., 2021). The paradigm consisted of a sampling phase and a testing phase, separated by a 15-second delay period in a closed-off start box. During the sampling phase, mice were forced to run a particular arm of the maze by the application of a barrier closing off the other reward arm. In the following test phase, mice were allowed to choose either arm and reward was only delivered for choosing the opposing arm to the preceding sample phase. Each mouse received five, 15-minute sessions of pre-training on the DNMP paradigm with gradually increasing delay in the week prior to surgery. Five additional training sessions of the same length, with full 15-second delay, were delivered after surgical recovery. All of these initial training sessions were carried out in the training maze, Context T: a grey, wooden, rectangular T-maze (66 cm long x 31 cm wide x 19 cm high), with opaque walls forming the central stem and different wall cues on each reward arm.

**Figure 1.**
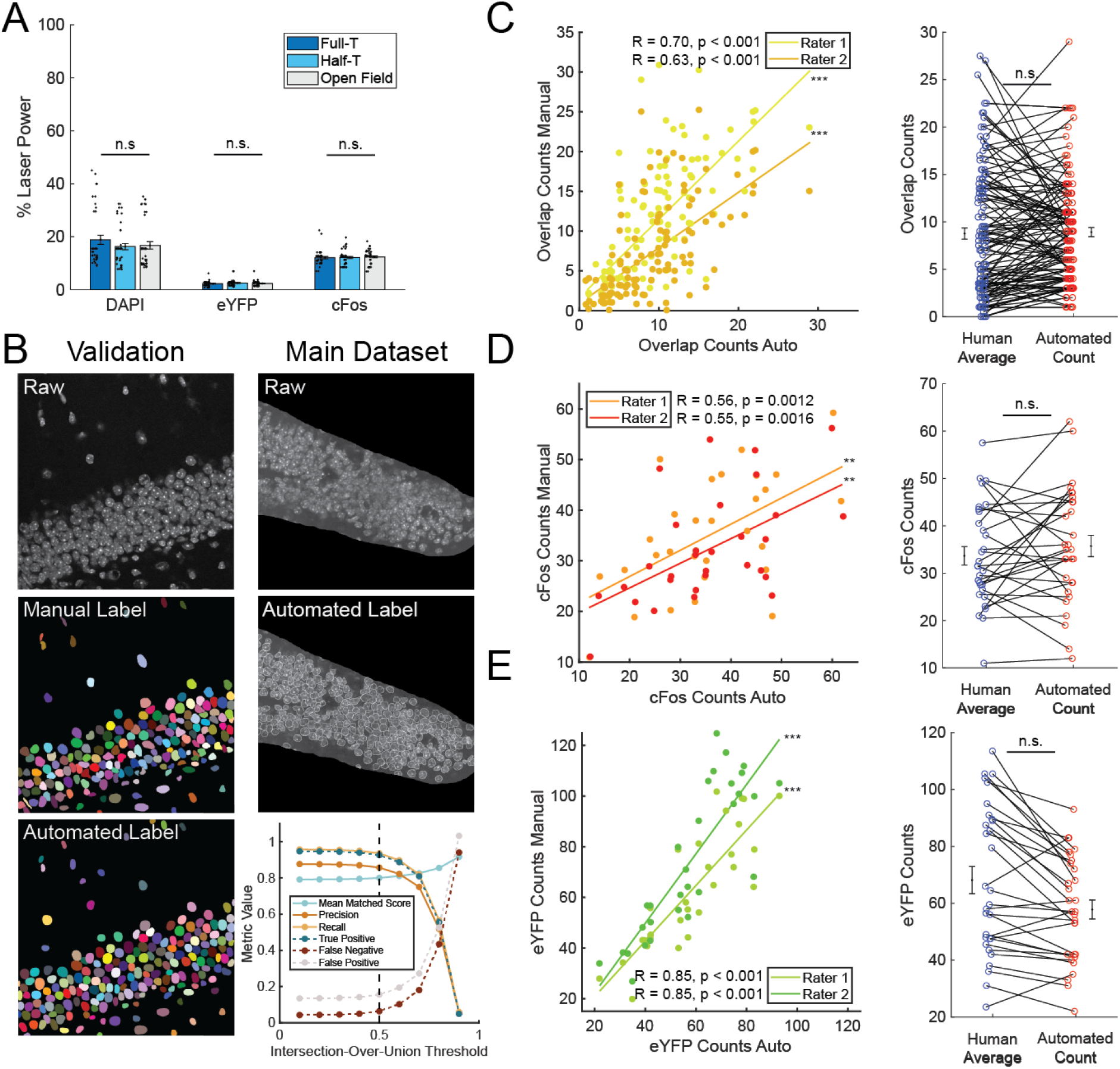
Similarity of cell counts supports the use of automated counting methods. **A)** Comparison of average laser power used during image acquisition across groups. No difference in power was found in the DAPI (F_(2,122)_ = 0.83, p = 0.44), eYFP (F_(2,122)_ = 0.66, p = 0.52) or cFos (F_(2,122)_ = 0,21, p = 0.81) channels, n = 40 images for Full-T and Half-T, 45 images for Open Field. **B)** Automated DAPI cell counts. Raw, manual label and automated label example from the training validation set. Colors for visualization purpose only. Raw and automated label example from the main data set. Bottom right: model validation metrics demonstrating high precision, accuracy, and matching to ground truth labels over a range of matching thresholds. Dashed black line indicates threshold used in dataset. True Positive, Negative, and False Negative rates shown normalized to ground truth. **C)** Left: Automated overlap counts strongly correlate with human raters. Pearson’s r and p values reported in figure. Right: no difference was found between average overlap counts of human raters and automated counting methods (t = -0.19, p = 0.85). n = 125 images. **D)** Left: Same as C but for cFos counts. Right: No difference in human or automated counting of cFos (t = -0.70, p = 0.47). n = 30 images. **E)** Left: Same as D but for eYFP counts. Right: No difference in human or automated counting of eYFP (t = 1.79, p = 0.079). Error bars represent +/- SEM. Scatter plots denote individual image samples jittered for clarity.

During the tagging window, mice were taken off dox for 48 hours and subsequently performed the DNMP paradigm in a novel T-maze, referred to as Context A: a beige, triangular, linoleum-lined T-maze (78 cm long x 78 cm wide x 18 cm high) with novel cue cards on the walls of the reward arms. The maze location, odor and floor texture were also changed relative to Context T. Context A had no walls on the stem, allowing mice to see the entire arena even when reward arm barriers were present. All mice were exposed to Context A for 20 minutes, with a minimum of ten DNMP trials performed. Immediately after tagging, the mice were placed back on dox for the remainder of the study. The following day, mice were pseudo-randomly split into three groups, counterbalancing for sex. The first group ran the full T-maze in Context A under the DNMP paradigm discussed above, for 20 minutes. The second group ran for 20 minutes on a one-sided route inside the same physical maze and at the same room position as Context A, but with the other arm blocked by opaque barriers (Fig. 2C, Context B). The mice in this group were evenly split between the left and right Half-T routes via pseudorandom assignment and were counterbalanced by sex. The third group was placed in a novel arena at the same room position as Contexts A and B and explored for 20 minutes without reward. This Open Field Context C was an empty, opaque gray box (41.5 cm long x 39.5 cm wide x 28.5 cm high) with no top.

**Figure 2.**
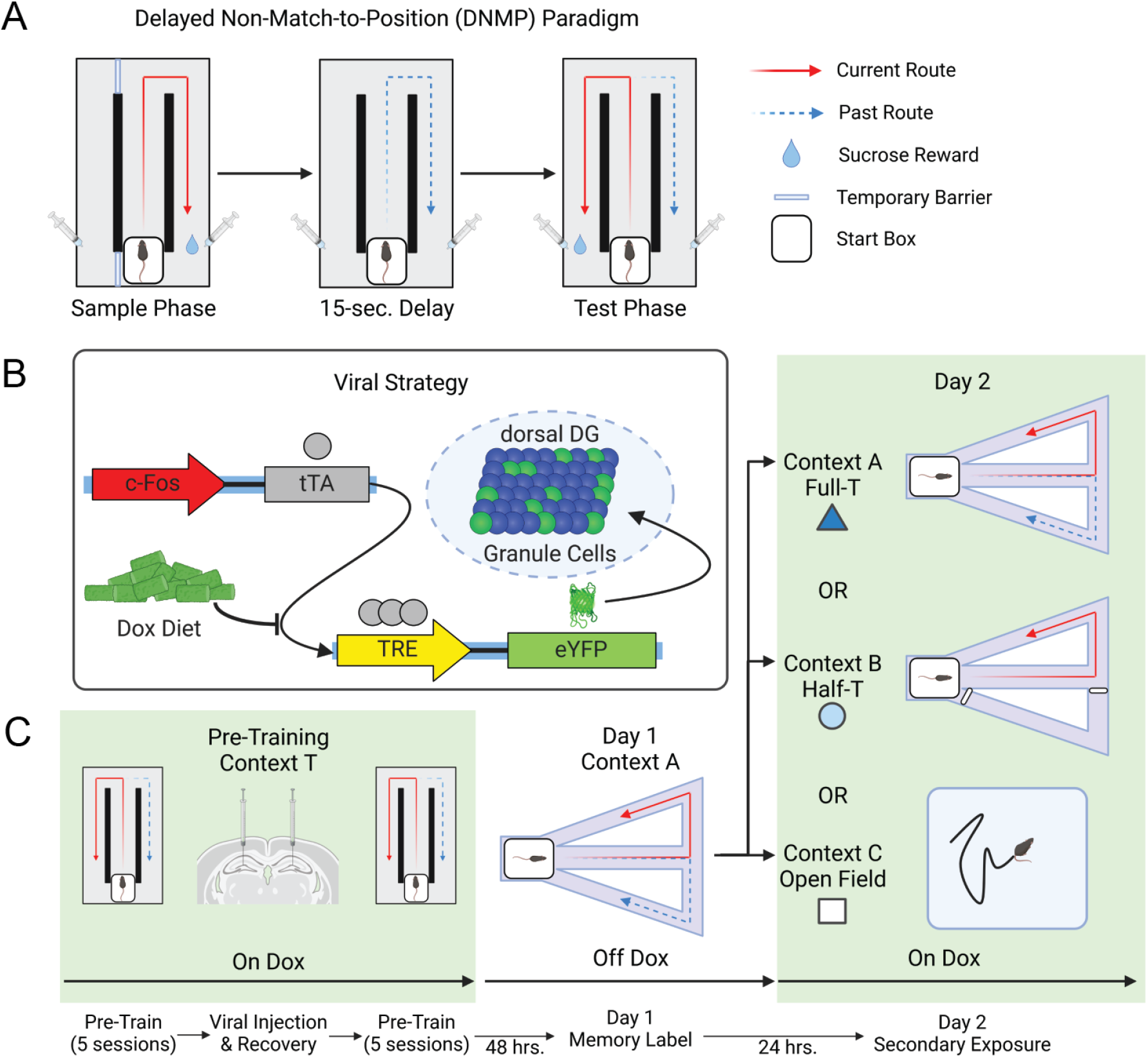
Behavioral and viral tagging paradigm used to visualize the spatial engram. **A)** Schematic representation of the delayed-non-match-to-position (DNMP) T-maze task. In the sample phase, an opaque barrier is inserted to force the mouse to traverse one route and receive sucrose reward. After a 15-second start box delay, the mouse must choose to traverse the opposite route to receive a second reward in the test phase. **B)** Viral constructs AAV9-cFos-tTA and AAV9-TRE-eYFP used in the engram labeling system. Endogenously expressed cFos in transfected Dentate Gyrus cells drives tetracycline-transactivator (tTA) which binds to the tetracycline response element (TRE) and drives expression of eYFP in the absence of doxycycline (Dox). **C)** Behavioral timeline. Mice (n=25) were pre-trained for 10 sessions, 5 before and 5 after surgical injection of the viral constructs outlined in B. Dox diet was removed from the home cage to open a memory labeling window. Day 1 performance of the DNMP-task in a novel Context A for 20 minutes caused cFos Dentate Gyrus cells to express eYFP. On Day 2 mice were exposed for 20 minutes to the full maze task (Full-T, Context A, n=8), the same maze but with one arm blocked (Half-T, Context B, n=8), or a novel arena (Open Field, Context C, n=9). Brains were collected 90 minutes after the behavioral experience on Day 2.

Deeplabcut (Mathis et al., 2018) was used to extract mouse position during behavioral trials from 50fps video recorded on an overhead Mako G-131c GigE camera (Allied Vision). Video timestamps, spatially scaled position, distance, and velocity information were extracted using the CMBHome framework (https://github.com/hasselmonians/CMBHOME/wiki).

### 2.5 Immunohistochemistry

Mice were euthanized 90 minutes after final behavior on Day 2 by administration of Euthasol (390 mg/kg) and anesthetized with Isoflurane prior to transcardial perfusion with saline and 10% formalin. Extracted brains were kept in formalin for 48 to 72 hours at 4 °C and transferred to 30% sucrose solution for approximately 72 hours at 4 °C to undergo cryoprotection. Cohorts included tissue from animals in all three groups, such that fixation and cryoprotection times were matched across groups. Brains were sliced using a cryostat into 50μm slices, and blocked for 2 hours at 4 °C in 1x phosphate-buffered-saline + 2% Triton (PBS-T) and 5% normal goat serum (NGS). Consistent with prior studies (Chen et al., 2019) slices were incubated for 48 hours at 4 °C with primary antibodies diluted in 5% NGS in PBS-T as follows: rabbit anti-cFos (1:1000, Abcam, #190289) and chicken anti-GFP (1:1000, ThermoFisher, #A10262). Subsequently, the slices were washed three times for 10 minutes in PBS-T, followed by a 2 hour incubation in the secondary antibodies diluted in 5% NGS in PBS-T as follows: Alexa 555 goat anti-rabbit (1:200; ThermoFihser, #A21429) and Alexa 488 goat anti-chicken (1:200, ThermoFisher, #A11039). Finally, the slices were mounted on slides using VECTASHIELD® Hardset™ Antifade Mounting Medium with DAPI (Vector Labs, #H-1500) and sealed with nail polish.

### 2.6 Image Acquisition

Images were acquired with an FV10i confocal laser-scanning microscope, using 60x magnification / 1.4 NA oil-immersion objective. For each image, three z-slices, separated by approximately 19.5μm, were acquired. The first z-slice was positioned at the lowest in-focus portion of the sample, the second was positioned at the center of the sample and the first was positioned at the highest in-focus portion of the imaged sample. The depth between the slices has been adjusted accordingly to ensure this composition, but was never set below 17 μm to minimize the potential of imaging the same cell in different z-planes. DAPI was acquired at 405nm with average laser power of 17.4% (SEM 1.43%), eYFP at 473nm with average laser power of 2.38% (SEM 0.17%) and cFos at 559nm with average laser power of 12.21% (SEM 0.40%). Before acquiring each image, the laser power of each channel (R, G, B) was configured to yield approximately equivalent intensity between all slices. To ensure images were collected with similar parameters across groups, we tested the laser power of each image on each channel across groups and found no significant differences in laser power between groups (Fig. 1A). Each image was acquired as a series of 1024×1024 pixel tiles which were subsequently stitched together to produce the final image (https://imagej.net/plugins/image-stitching). Only one set of stitching parameters was used, ensuring any stitching artefacts were consistent across groups. The images used 16 bit-depth encoding and all image processing was handled in ImageJ / FIJI (https://imagej.nih.gov/ij/) with the Bioformats toolbox (https://imagej.net/formats/bio-formats).

### 2.7 Cell Counting

Image acquisition was performed only on slices with successful targeting to the dorsal DG. cFos+ and eYFP+ cells in the upper and lower DG blades were counted using five 50um coronal slices of dorsal DG in each animal. Cell counts from each of the three z-planes were summed for that slice and averaged across slices for each mouse. DAPI was counted using a StarDist neural network trained and validated on a subset of the data (Fig 1B,Schmidt et al., 2018; Weigert et al., 2020). Segmentation metrics (precision, recall, and matching to ground truth labels) confirmed model performance over a range of label matching thresholds. An intersection-over-union threshold of 0.5% was used as a tradeoff between match score and other metrics. The number of cFos+ and eYFP+ cells in the DG region was quantified using an automated ImageJ pipeline that carried out iterative thresholding followed by exclusion of non-circular objects (https://mcib3d.frama.io/3d-suite-imagej/plugins/Segmentation/3D-Iterative-Segmentation/). Parameters were calibrated to include cells even where stitched images showed intensity variability. In order to quantify the number of overlapping cFos+ and eYFP+ cells, an automated algorithm was used to carry out pairwise comparisons between the pixels of each eYFP and cFos cell and the results were filtered to only include overlapping cells of a comparable size (within at least 50% of each-other’s size) that were mostly overlapping (85% of smaller object). The accuracy of the automatic counter was verified against a manually scored dataset created by two experimenters blind to the experimental conditions (Fig. 1C-E). Automated counts were used for increased reproducibility. Statistical chance for overlap was calculated as 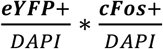 and compared against the counted overlaps normalized to the whole dentate population as 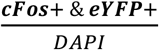.

### 2.8 Statistical Analysis

Bar graphs are reported as means +/- SEM. One way and mixed effects ANOVA tests were used to assess group differences for both cell counts and behavior, and follow-up comparisons made where appropriate using independent T-tests (Tukey’s HSD). Repeated measures comparisons including overlap against statistical chance were performed using paired t-tests while all other comparisons were made with independent t-tests. Bonferroni-Holm corrected p-values are reported where multiple t-test comparisons are made. Correlations were run using Pearson’s R. Two mice were excluded (1 Full-T, 1 Half-T) from velocity and distance comparisons (Fig. 4C,D) on the basis of corrupted video files but were otherwise included for all other tests. Test statistics, groups sizes, and p-values are reported in figure legends. All tests were performed in Matlab using publicly available functions. For all figures, * = p < 0.05, ** = p < 0.01, *** = p < 0.001.

### 2.9 Data Availability

All relevant data supporting the findings of this study are available from the corresponding author upon reasonable request.

## 3. RESULTS

To investigate cells in Dentate Gyrus (DG) associated with spatial navigation memories, we trained mice on a delayed-non-match-to-position (DNMP) task (Fig. 2A). A dox-inducible viral labeling strategy was used to selectively label cFos*+* DG cells associated with learning (Fig. 2B). After pre-training and surgical recovery, dox diet was removed from the cage and mice were exposed to a novel T-maze (Context A) in which they performed the DNMP task for 20 minutes (Fig. 2C). The mice were returned to dox to prevent off-target labeling and exposed on the following day to another environment in the same position with respect to room cues as Context A. The first group (Full-T) repeated the DNMP behavior in Context A as before. The second group was returned to the same physical arena as Context A, but with one arm permanently blocked (Context B). This arena was otherwise similar with respect to timing delays and sucrose rewards and allowed mice visual access to the other side of the maze. The final group was exposed to a novel open field arena without reward contingency (Context C).

We first examined the size and degree of overlap between DG cell ensembles to determine the level of representational similarity across days in each group. Immunohistochemical staining revealed two populations, the first from the Day 1 experience (eYFP) and the second from the Day 2 experience (cFos) with partial reactivation (overlap; Fig. 3A-F). Performance of the DNMP T-maze task on Day 1 yielded similar ensemble size across groups as expected (Fig. 4A). Interestingly, no difference was found in Day 2 ensemble size across groups despite the Open Field group experiencing a novel environment.

**Figure 3.**
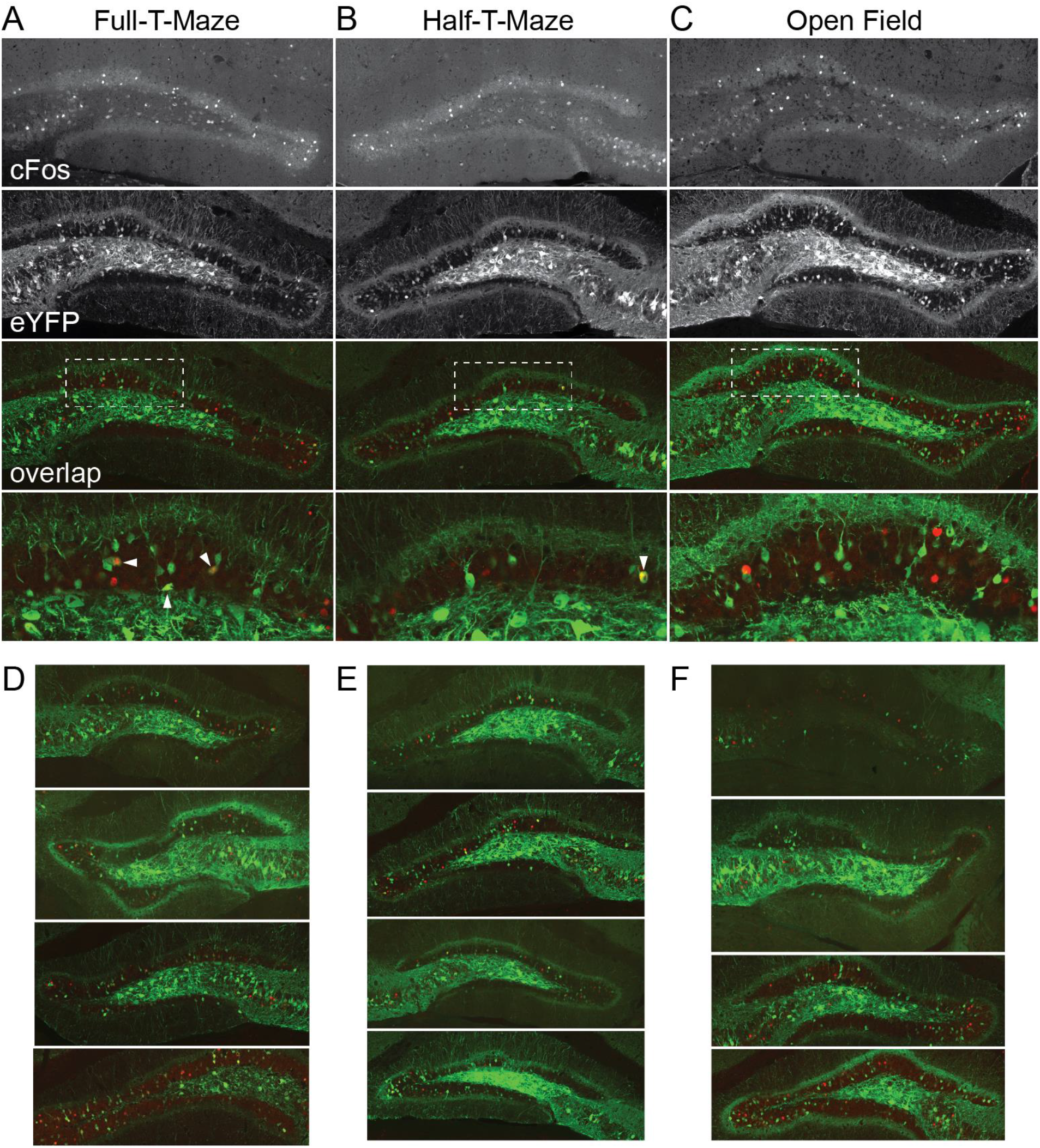
Histological staining reveals distinct, partially overlapping spatial engram populations in DG. **A-C)** Representative 60x confocal images of DG from the Full-T, Half-T and Open Field groups. Top row: cFos signal. Upper middle row: eYFP signal. Lower middle row: merge of the above rows (cFos red, eYFP green). Bottom: Zoomed section from the above merge indicated by the white dashed box. White triangles indicate overlap cells active on both days. **D-F)** Further examples from four different animals per group outlined above in A-C.

Next, we normalized the number of overlapping cells to the whole dentate population and compared each group to statistical chance. This allowed testing of the primary hypothesis of the experiment as summarized in Figure 4B. Both the Full-T and Half-T exposures resulted in significantly overlapping populations, while the Open Field exposure did not (Fig. 4B). Additionally, we found a dissociation of reactivation level across groups, with the Full-T experiencing highest overlap and Open Field experiencing the least (Fig. 4B). These results suggest an effect of both novelty and trajectory experience on the recruitment and reactivation of the DG spatial engram ensemble.

**Figure 4.**
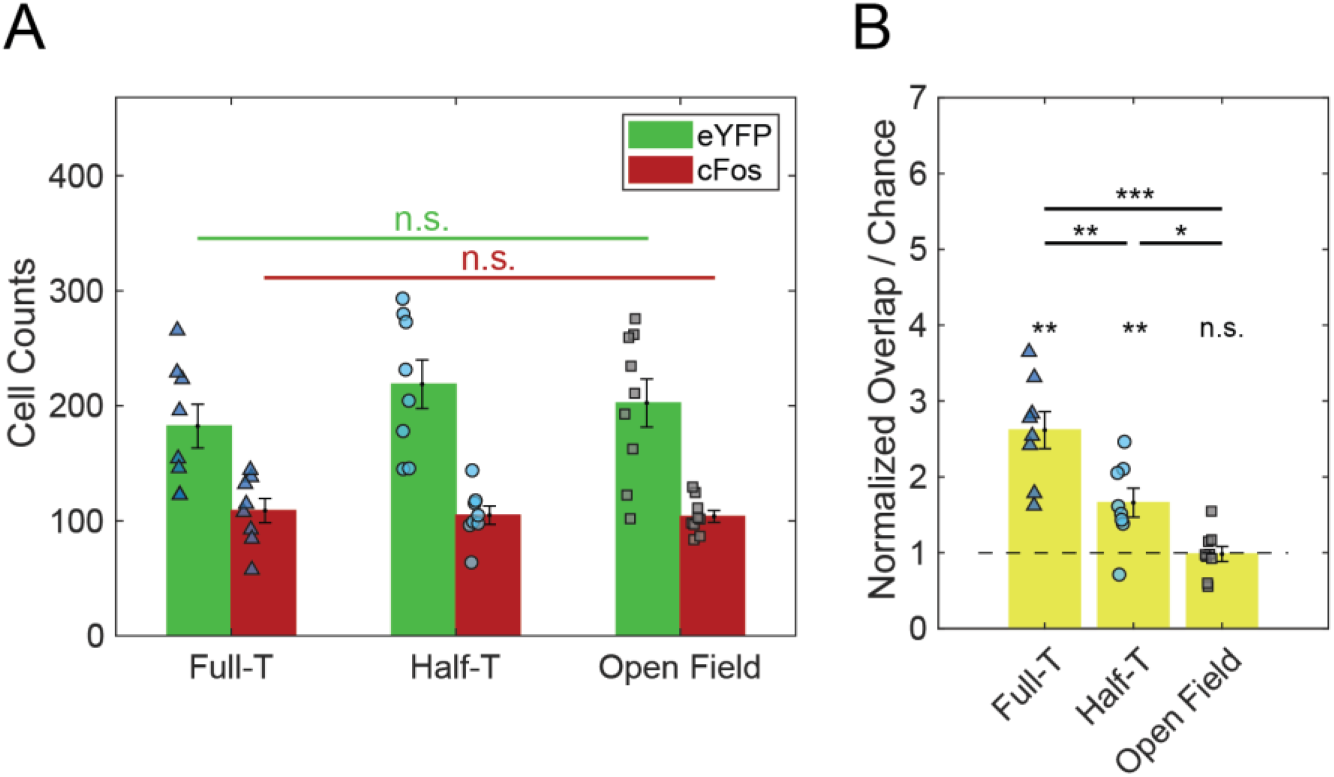
DG encodes different routes within a maze using partially non-overlapping populations. **A)** There was no difference in Day 1 ensemble size (F_(2,22)_ = 76, p = 0.48) or in Day 2 ensemble size (F_(2,22)_ = 0.11, p = 0.90) across groups. **B)** Experience of the Day 2 environments and tasks resulted in ensemble reactivation (overlap) above statistical chance in the Full-T (t = 6.23, p = 0.0012) and Half-T (t = 4.09, p = 0.0093), but not the Open Field groups (t = -1.04, p = 0.33). Additionally, normalized reactivation in all groups differed from one another (F_(2,22)_ = 20.44, p < 0.001). Post-hoc comparisons revealed significant differences between all groups (Tukey’s HSD; Full-T vs Half-T: p = 0.0040; Full-T vs Open Field: p < 0.001; Half-T vs Open Field: p = 0.038) Error bars represent +/- SEM. Scatter plots denote individual subjects (triangle = Full-T, circle = Half-T, square = Open Field). N = 8 mice in Full-T and Half-T groups, 9 mice in Open Field.

To rule out the possibility of confounding effects such as task engagement, laps traversed, and behavioral activity level on engram ensemble recruitment, we analyzed behavior across groups. All groups performed the task above chance nor did we find a difference in accuracy across groups on Day 1 (Fig. 5A). There was no change in accuracy across days for the Full-T group. These data demonstrate that the mice grasped the task and transferred learning successfully from training Context T to the novel Context A. Next, we examined the number of laps, defined as a trajectory from the start box to the reward then back to the start box, across groups and days. While the Full-T and Half-T groups did not differ in Day 2 laps, the Half-T group performed significantly fewer laps on Day 2 compared to Day 1 baseline (Fig. 5B), possibly reflecting a shift to a different behavioral demand (Satvat et al., 2011). For a more general comparison of behavior, we analyzed total distance traveled and average running speed in all groups across days. We found a main effect of day but no effect of group or interaction in either metric (Fig. 5C,D). Anecdotally, we observed mice in the Half-T condition on Day 2 actively investigating the barrier at the choice point which could explain the reduced lap number in some animals even while the distance traveled and average running speed remained constant across groups on Day 2. To follow up this result, we tested for possible correlations between ensemble reactivation and behavior. No correlation was found between the normalized ensemble overlap and any behavioral metric (task accuracy, number of laps, distance, or average speed) on Day 1 or Day 2 in any group (Fig. 6A-D). These analyses suggest that the degree of engram reactivation is not a result of trajectory length or self-motion cues, but is instead related to behavioral contingencies and experienced trajectories, as hypothesized.

**Figure 5.**
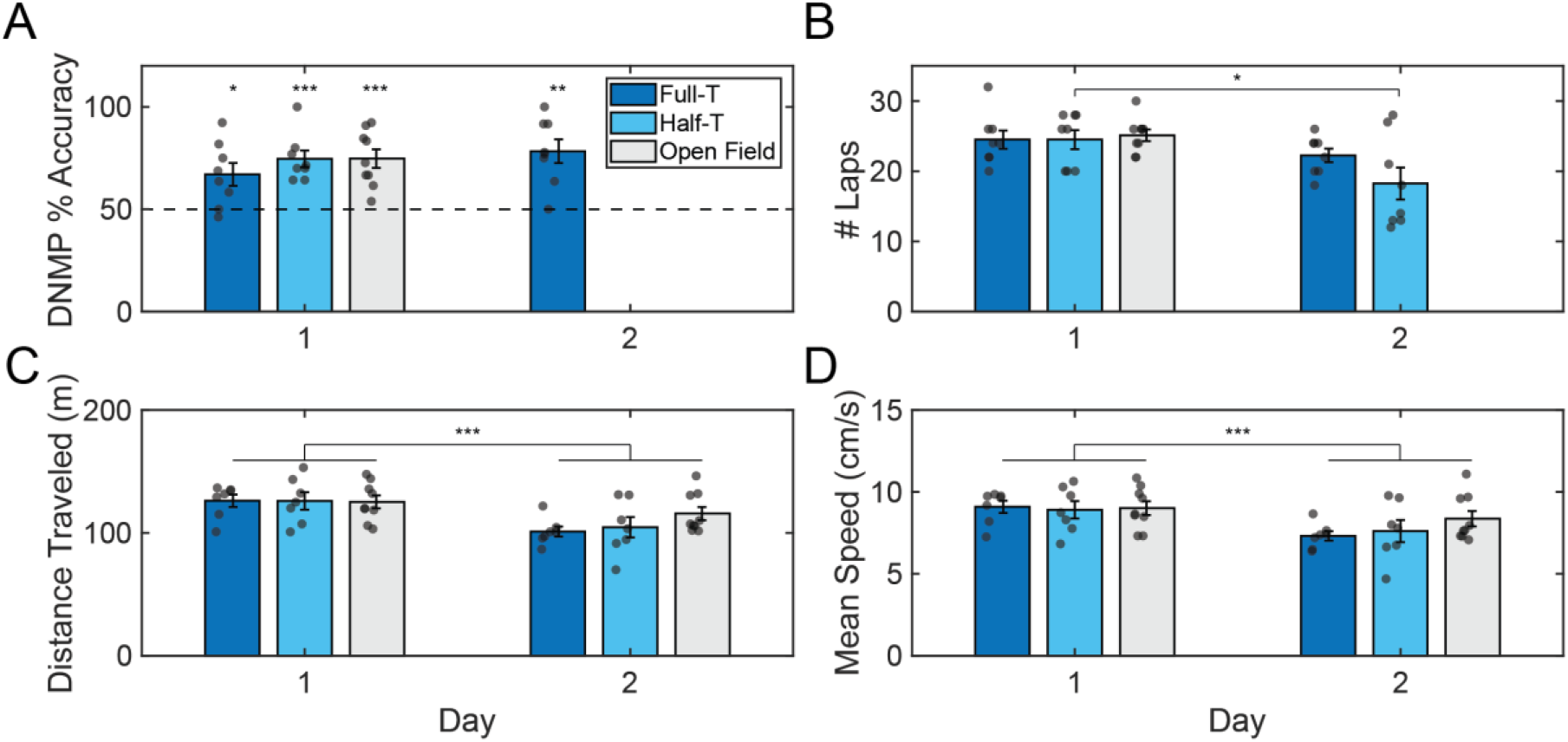
DG spatial engram composition relates to experienced trajectories not distance or velocity. **A)** All groups performed above chance (One-sample t-test; Day 1: Full-T t = 3.05, p = 0.018; Half-T t = 5.99, p = 0.0022; Open Field t = 5,44, p = 0.0018; Day 2: Full-T t = 4.92, p = 0.0034). No differences in DNMP task accuracy were found across groups on Day 1 (F_(2,22)_ = 0.85, p = 0.44)) nor did the Full-T group perform differently across days (t = -1.78, p = 0.12). **B)** Groups did not run different numbers of laps on Day 1 (F_(2,22)_ = 0.097, p = 0.91), nor did the Half-T group run fewer laps relative to the Full-T group on Day 2 (t = 1.67, p = 0.14). The Full-T group ran similar laps across days (t = 1.94, p = 0.19) but the Half-T ran fewer on Day 2 relative to Day 1 (t = 3.31, p = 0.039). **C)**The total distance traveled did not differ across groups (F_(2,20)_ = 0.47, p = 0.63) but was lower on Day 2 (F_(1,20)_ = 41.11, p < 0.001) with no interaction effect (F_(2,20)_ = 2.82, p = 0.084). **D)** The average velocity did not differ across groups on Day 1 (F_(2,20)_ = 0.43, p = 0.66) but was lower on Day 2 (F_(1,20)_ = 24.02, p < 0.001) with no interaction effect (F_(2,20)_ = 1.77, p = 0.20). Error bars represent +/- SEM. Scatter plots denote individual mice. N = 9 mice in the Open Field group. For accuracy and lap comparisons, N = 8 mice in Full-T and Half-T groups. N = 7 mice in Full-T and Half-T groups for velocity and distance comparisons.

**Figure 6.**
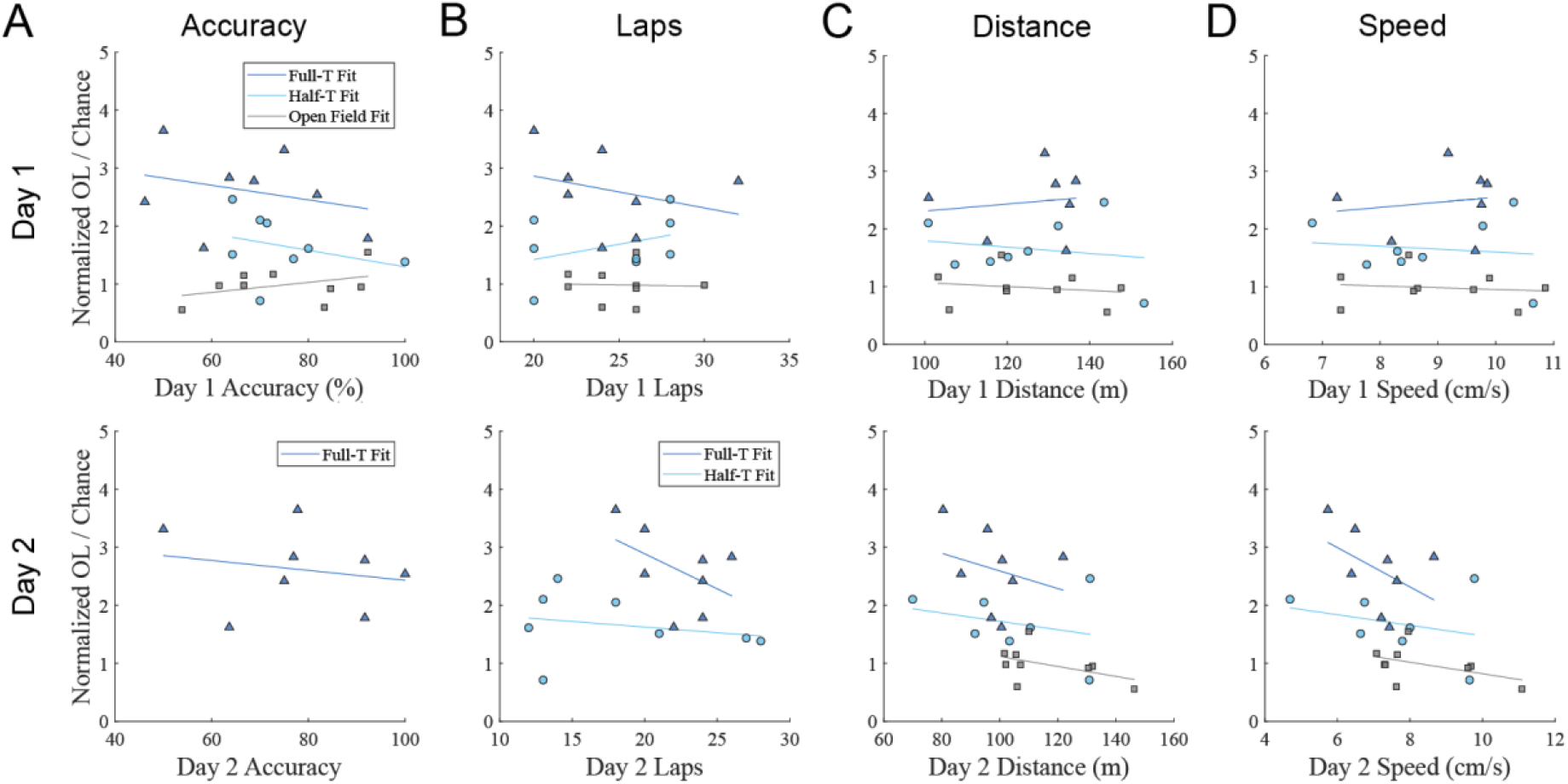
Engram reactivation does not correlate to behavioral outcomes. **A)** Top: No relationship was found between Day 1 DNMP accuracy and degree of engram reactivation in any group (Full-T: r = -0.29, p = 0.49; Half-T: r = -0.31, p = 0.92; Open Field: r = 0.39, p = 0.88). Bottom: Same as above but for Day 2, Full-T (r = -0.20, p = 0.63). **B)** Same as A but for number of laps on Day 1 (Full-T: r = -0.29, p = 0.97; Half-T: r = 0.37, p = 1.00; Open Field: r = -0.03, p = 0.93) and Day 2 (Full-T: r = -0.47, p = 0.47; Half-T: r = -0.23, p = 0.58) **C)** Same as B but for distance traveled on Day 1 (Full-T: r = 0.14, p = 0.77; Half-T: r = -0.18, p = 1,00; Open Field: r = -0.18, p = 1.00) and Day 2 (Full-T: r = -0.27, p = 1.00; Half-T: r = -0.28, p = 0.55; Open Field: r = -0.13, p = 0.62) **D)** Same as C but for distance traveled on Day 1 (Full-T: r = 0.15, p = 1.00; Half-T: r = -0.12, p = 0.77; Open Field: r = -0.13, p = 1.00) and Day 2 (Full-T: r = -0.44, p = 0.55; Half-T: r = -0.28, p = 0.54; Open Field: r = -0.47, p = 0.62) N = 9 mice in the Open Field group. For accuracy and lap comparisons, N = 8 mice in Full-T and Half-T groups. N = 7 mice in Full-T and Half-T groups for velocity and distance comparisons.

Finally, we examined our data broken down by subject sex and direction of the maze experienced in the Half-T group. While each group lacked sufficient number of subjects for rigorous comparison by sex within groups, observationally we saw little difference in average Day 1 or Day 2 ensemble size or in normalized engram reactivation (Fig. 7A). At the behavioral level, male and female mice exhibited little difference in any behavioral metric on either day (Fig. 7B). In the Half-T group, we observed minimal difference between the Left-T and Right-T subgroups at either the ensemble level (Fig. 7C) or the behavioral level (Fig. 7D). These similarities were used as grounds to combine animals by sex and by Half-T subgroup in the previous analyses, although follow up studies with larger group sizes would be needed to confirm the lack of effect.

**Figure 7.**
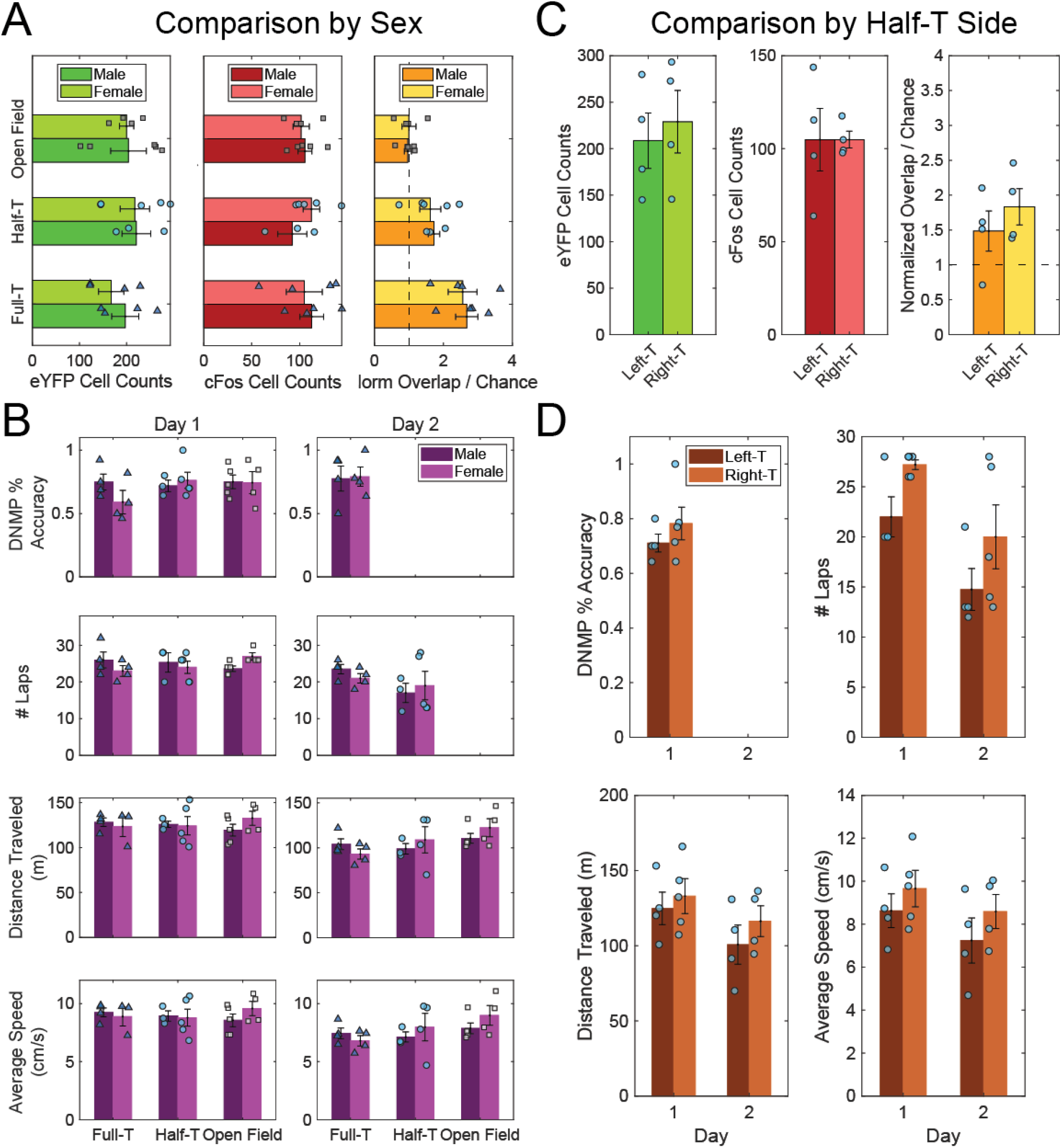
Engram composition and behavior does not appear to differ across sex or experienced trajectory in the Half-T group. **A)** The size of the Day 1 and Day 2 ensembles, as well as degree of reactivation (overlap cells) appears similar between male and female mice of all groups. **B)** Only minor sex differences were observed for any behavioral metric on either Day 1 (Left) or Day 2 (right) for all groups. **C)** Same as A but comparing the Left-T and Right-T subgroups in the Half-T group, similarly little difference observed. **D)** Only minor subgroup differences were observed for behavioral metrics on Day 1 and Day 2. Error bars represent +/- SEM. Scatter plots denote individual mice.

## 4. DISCUSSION

We trained mice to run a delayed-non-match-to-position (DNMP) T-maze task and used an activity-dependent viral labeling strategy to visualize cell populations from different days associated with goal-directed navigation, i.e. spatial engrams. We found that repeated experience with the same two-route spatial working memory task and physical location across days yielded the highest degree of engram similarity (Full-T; Fig. 4B). Mice performing a navigation task in the same physical arena and room position but with only one route (Half-T) showed higher reactivation of the original population than chance levels, but less than the Full-T group (Fig. 4B). The Open Field group exhibited the least overlap, no different than chance (Open Field, Fig. 4B). In addition, we observed no difference in the size of the Day 2 ensemble despite differences in memory demand and novelty based on the experienced arena (Fig. 4A). Together, these results are consistent with past studies implicating the DG in spatial memory processing (McNaughton et al., 1989; Emerich & Walsh, 1990; Xavier et al., 1999), including T-maze tasks with long delays (Emerich & Walsh, 1989; Costa et al., 2005) and build on previous applications of immediate early genes (IEGs) to study the overlap of populations involved in hippocampal spatial and task-specific memories (Guzowski et al., 1999; Satvat et al., 2011).

### 4.1 Spatial vs fear engrams

We set out to test whether methodologies typically used to label contextual memories in dorsal DG could be used to probe aspects of navigational memory for specific routes within a larger environment. Previous studies of memory ensemble, or engram, composition have largely focused on the formation and reactivation of contextual memories with strong emotional valence, in particular fear (Liu et al., 2012b; Ramirez et al., 2013; Redondo et al., 2014; Chen et al., 2019; Sun et al., 2020). However, physiological recordings from DG during navigation have revealed spatial preferences of DG granule and mossy cells (Jung & McNaughton, 1993; Leutgeb et al., 2007; Neunuebel & Knierim, 2012, 2014; GoodSmith et al., 2017, 2022) which may be stable across days (Hainmueller & Bartos, 2018; Cholvin et al., 2021). While IEGs have been used to examine cell population reactivation across or within open field navigation contexts (Guzowski et al., 1999; VanElzakker et al., 2008), this approach could not distinguish an engram ensemble coding for contextual cues (e.g. odor, distal visual landmarks, etc.) from one encoding specific navigational trajectories (e.g. serially activated place fields). In our study, the Half-T group had lower ensemble reactivation than the Full-T group, but higher reactivation than the Open Field group and statistical chance (Fig. 4B). Some contribution of pattern separation as a result of differences in behavioral demand between a spatial reference memory task (Full-T) and a sensory-guided navigation task with similar delays and rewards (Half-T) is possible (Satvat et al., 2011). However, we found largely no difference in behavior across groups except a small reduction in lap number across days in the Half-T group, potentially due to some animals actively investigating the barrier during laps (Fig. 4B). We found largely no correlation between engram reactivation and behavioral measures (Fig. 5) in agreement with past work showing little relationship between freezing and fear engram ensemble activity (Zaki et al., 2022) or locomotion and Fos levels in hippocampus (VanElzakker et al., 2008). Based on these finding and previous evidence for spatial specificity in DG granule cells, we hypothesize that different routes are encoded by different spatial engram populations.

The relationship between Fos expression, memory encoding, and spatial correlates of cell activity is complex. One recent study linked the degree of Fos expression to the reliability and stability place fields in CA1 during familiar task performance (Pettit et al., 2022). Interestingly, clusters of co-active cells with strong Fos expression exhibited place fields across large sections of the environment, consistent with prior work in replay and theta sequences suggesting that correlated cells chunk spatial information (Foster & Wilson, 2006; Johnson & Redish, 2007; Gupta et al., 2012). Conversely, a previous engram study using electrophysiology found that cFos tagged CA1 cells displayed strong contextual firing but poorer spatial stability than non-tagged cells (Tanaka et al., 2018). One possible explanation for the disparity is the level of experience with the environment, because cFos is both driven by firing activity and helps maintain spatial coding accuracy in existing place cells (Pettit et al., 2022). In our study, mice were exposed to novel environments during tagging, likely driving formation of new engram ensembles similar to Tanaka et al. However, the animals had extensive pre-training on the task itself and transferred learning across environments (Fig. 5A). It is therefore unsurprising that the Full-T group displayed reactivation above chance levels and above the Half-T group, indicating stability of the cFos tagged ensemble across days despite the novel environment, in line with the experienced animals and findings of Pettit et al. Follow up studies could compare ensembles tagged in the training context, or pre-trained versus naïve mice, to disambiguate the competing factors of pattern completion and pattern separation on spatial engram recruitment (Nakashiba et al., 2012; Santoro, 2013). Further, investigating the cellular dynamics in DG during task demand updating (as in the Half-T group on Day 2) would provide valuable insight into real-time feedback influencing memory and spatial-associated cell ensembles.

### 4.2 Size of the Engram Ensemble

cFos IEG expression arises from neural activity and leads to various plasticity-related changes within a cell (Labiner et al., 1993). Novelty is an important factor for inducing plasticity in the hippocampus, including DG (McNaughton & Morris, 1987; Kitchigina et al., 1997; Straube, Korz, & Frey, 2003; Straube, Korz, Balschun, et al., 2003; Davis et al., 2004). We observed a larger Day 1 ensemble relative to the Day 2 ensemble in all groups, consistent with a novelty effect in the Full-T and Half-T groups (Fig. 4A) and with prior work using similar techniques (Zaki et al., 2022). However, we made no formal comparison of this effect due to the difference in cFos detection between IHC and viral tagging methods and because animals ran less distance and at lower speed on the second day (Fig. 5C,D). Past work using alternate Fos detection methods similarly demonstrated increases in DG Fos expression after novel, but not familiar, environmental exploration (VanElzakker et al., 2008). Interestingly, the Open Field group underwent a novel context exposure on each day and exhibited a similar Day 2 ensemble size to the other groups, which suggests an impact not only of novelty but also task in this study (Fig. 4A). Given similar ensemble detection methods to the other groups, we might have expected the Open Field group to have larger average Day 2 ensemble size, but this was not the case. The open field free-exploration task has no route constraints by design, and these mice were just as active as their Full-T and Half-T counterparts (Fig. 5C,D). Thus, both the navigational memory demand and reward contingencies likely play a role in DG ensemble recruitment in addition to novelty alone (Costa et al., 2005).

### 4.3 Composition of sub-ensembles for pattern separation

One question concerns whether the DG contains cells with task-modulated spatial tuning, i.e. splitter cells, like those observed in CA1 (Wood et al., 2000; Ferbinteanu & Shapiro, 2003; Griffin et al., 2007; Kinsky et al., 2020). Splitter cells can emerge early in learning and may be modulated by the turn direction, task phase, or both on the DNMP T-maze task (Levy et al., 2021). DG place fields remap based on task engagement, hinting at the flexibility of DG cells within the same physical location (Shen et al., 2021). Interestingly, CA1 splitter cells appear more stable than classic place fields across days, which might help to explain increased overlap observed in the Full-T group relative to the Half-T group which had no chance to demonstrate splitter behavior (Fig 4B; Kinsky et al., 2020). Splitter cells may indeed serve an additional pattern separation function in DG for this task by discriminating otherwise similar stem trajectories and improving pattern completion of orthogonal sub-ensembles coding for separate reward arm routes (O’Reilly & McClelland, 1994; Wood et al., 2000; Ferbinteanu & Shapiro, 2003; Hasselmo & Eichenbaum, 2005; Nakashiba et al., 2012; Neunuebel & Knierim, 2012, 2014; GoodSmith et al., 2017, 2019; Senzai & Buzsáki, 2017; Hainmueller & Bartos, 2018).

### 4.4 Conclusion

We tested whether experience with specific navigation routes could be dissected in the dentate gyrus using engram tagging and visualization strategies. We found that repeated experience of a two-route maze task across days recruited a more similar ensemble than exposure to a novel open field arena or exposure to a one-route task within the same T-maze arena. The experimental design offers a means to study aspects of spatial navigation using traditional engram tagging techniques. Additionally, our results suggest the dentate gyrus performs its role in pattern separation and spatial navigation by the activation of partially non-overlapping sub-ensembles for different routes in a larger context.

## Acknowledgments

We thank the Hasselmo and Ramirez labs for thoughtful feedback and commentary on the work.

This work was supported by the National Institutes of Health [grant numbers: R01 MH052090, MH060013, MH120073, HD101402-02; and DP5 OD023106-01], and the Office of Naval Research [grant numbers: MURI N00014-16-1-2832, N00014-19-1-2571; and DURIP N00014-17-1-2304].

## Author Contributions

Conceptualization, L.K.W., S.R., and M.E.H.; Methodology, L.K.W. and I.K.; Investigation, L.K.W. and I.K.; Writing – Original Draft, L.K.W., I.K.; Writing – Review & Editing, L.K.W., I.K., S.R., and M.E.H.; Funding Acquisition, M.E.H. and S.R.; Formal Analysis, L.K.W. and I.K.; Visualization, L.K.W.; Supervision, M.E.H. and S.R.

## References

Chen, B. K., Murawski, N. J., Cincotta, C., McKissick, O., Finkelstein, A., Hamidi, A. B., Merfeld, E., Doucette, E., Grella, S. L., Shpokayte, M., Zaki, Y., Fortin, A., & Ramirez, S. (2019). Artificially Enhancing and Suppressing Hippocampus-Mediated Memories. Current Biology, 29(11), 1885-1894.e4. https://doi.org/10.1016/j.cub.2019.04.065

Cholvin, T., Hainmueller, T., & Bartos, M. (2021). The hippocampus converts dynamic entorhinal inputs into stable spatial maps. Neuron, 109(19), 3135-3148.e7. https://doi.org/10.1016/j.neuron.2021.09.019

Costa, V. C. I., Bueno, J. L. O., & Xavier, G. F. (2005). Dentate gyrus-selective colchicine lesion and performance in temporal and spatial tasks. Behavioural Brain Research, 160(2), 286–303. https://doi.org/10.1016/j.bbr.2004.12.011

Davis, C. D., Jones, F. L., & Derrick, B. E. (2004). Novel Environments Enhance the Induction and Maintenance of Long-Term Potentiation in the Dentate Gyrus. The Journal of Neuroscience, 24(29), 6497–6506. https://doi.org/10.1523/JNEUROSCI.4970-03.2004

Denny, C. A., Kheirbek, M. A., Alba, E. L., Tanaka, K. F., Brachman, R. A., Laughman, K. B., Tomm, N. K., Turi, G. F., Losonczy, A., & Hen, R. (2014). Hippocampal Memory Traces Are Differentially Modulated by Experience, Time, and Adult Neurogenesis. Neuron, 83(1), 189–201. https://doi.org/10.1016/j.neuron.2014.05.018

Emerich, D. F., & Walsh, T. J. (1989). Selective working memory impairments following intradentate injection of colchicine: Attenuation of the behavioral but not the neuropathological effects by gangliosides GM1 and AGF2. Physiology & Behavior, 45(1), 93–101. https://doi.org/10.1016/0031-9384(89)90170-4

Emerich, D. F., & Walsh, T. J. (1990). Cholinergic cell loss and cognitive impairments following intraventricular or intradentate injection of colchicine. Brain Research, 517(1–2), 157–167. https://doi.org/10.1016/0006-8993(90)91021-8

Ferbinteanu, J., & Shapiro, M. L. (2003). Prospective and Retrospective Memory Coding in the Hippocampus. Neuron, 40(6), 1227–1239. https://doi.org/10.1016/S0896-6273(03)00752-9

Foster, D. J., & Wilson, M. A. (2006). Reverse replay of behavioural sequences in hippocampal place cells during the awake state. Nature, 440(7084), Article 7084. https://doi.org/10.1038/nature04587

GoodSmith, D., Chen, X., Wang, C., Kim, S. H., Song, H., Burgalossi, A., Christian, K. M., & Knierim, J. J. (2017). Spatial Representations of Granule Cells and Mossy Cells of the Dentate Gyrus. Neuron, 93(3), 677-690.e5. https://doi.org/10.1016/j.neuron.2016.12.026

GoodSmith, D., Kim, S. H., Puliyadi, V., Ming, G., Song, H., Knierim, J. J., & Christian, K. M. (2022). Flexible encoding of objects and space in single cells of the dentate gyrus. Current Biology, 0(0). https://doi.org/10.1016/j.cub.2022.01.023

GoodSmith, D., Lee, H., Neunuebel, J. P., Song, H., & Knierim, J. J. (2019). Dentate Gyrus Mossy Cells Share a Role in Pattern Separation with Dentate Granule Cells and Proximal CA3 Pyramidal Cells. The Journal of Neuroscience, 39(48), 9570–9584. https://doi.org/10.1523/JNEUROSCI.0940-19.2019

Griffin, A. L., Eichenbaum, H., & Hasselmo, M. E. (2007). Spatial Representations of Hippocampal CA1 Neurons Are Modulated by Behavioral Context in a Hippocampus-Dependent Memory Task. Journal of Neuroscience, 27(9), 2416–2423. https://doi.org/10.1523/JNEUROSCI.4083-06.2007

Gupta, A. S., van der Meer, M. A. A., Touretzky, D. S., & Redish, A. D. (2012). Segmentation of spatial experience by hippocampal theta sequences. Nature Neuroscience, 15(7), 1032–1039. https://doi.org/10.1038/nn.3138

Guzowski, J. F., McNaughton, B. L., Barnes, C. A., & Worley, P. F. (1999). Environment-specific expression of the immediate-early gene Arc in hippocampal neuronal ensembles. Nature Neuroscience, 2(12), 1120–1124. https://doi.org/10.1038/16046

Hainmueller, T., & Bartos, M. (2018). Parallel emergence of stable and dynamic memory engrams in the hippocampus. Nature, 558(7709), 292–296. https://doi.org/10.1038/s41586-018-0191-2

Hasselmo, M. E., & Eichenbaum, H. (2005). Hippocampal mechanisms for the context-dependent retrieval of episodes. Neural Networks, 18(9), 1172–1190. https://doi.org/10.1016/j.neunet.2005.08.007

Hasselmo, M. E., & Wyble, B. P. (1997). Free recall and recognition in a network model of the hippocampus: Simulating effects of scopolamine on human memory function. Behavioural Brain Research, 89(1–2), 1–34. https://doi.org/10.1016/S0166-4328(97)00048-X

Johnson, A., & Redish, A. D. (2007). Neural Ensembles in CA3 Transiently Encode Paths Forward of the Animal at a Decision Point. Journal of Neuroscience, 27(45), 12176–12189. https://doi.org/10.1523/JNEUROSCI.3761-07.2007

Jung, M. W., & McNaughton, B. L. (1993). Spatial selectivity of unit activity in the hippocampal granular layer. Hippocampus, 3(2), 165–182. https://doi.org/10.1002/hipo.450030209

Kinsky, N. R., Mau, W., Sullivan, D. W., Levy, S. J., Ruesch, E. A., & Hasselmo, M. E. (2020). Trajectory-modulated hippocampal neurons persist throughout memory-guided navigation. Nature Communications, 11, 2443. https://doi.org/10.1038/s41467-020-16226-4

Kitamura, T., Ogawa, S. K., Roy, D. S., Okuyama, T., Morrissey, M. D., Smith, L. M., Redondo, R. L., & Tonegawa, S. (2017). Engrams and circuits crucial for systems consolidation of a memory. Science, 356(6333), 73–78. https://doi.org/10.1126/science.aam6808

Kitchigina, V., Vankov, A., Harley, C., & Sara, S. J. (1997). Novelty-elicited, Noradrenaline-dependent Enhancement of Excitability in the Dentate Gyrus. European Journal of Neuroscience, 9(1), 41–47. https://doi.org/10.1111/j.1460-9568.1997.tb01351.x

Labiner, D., Butler, L., Cao, Z., Hosford, D., Shin, C., & McNamara, J. (1993). Induction of c-fos mRNA by kindled seizures: Complex relationship with neuronal burst firing. The Journal of Neuroscience, 13(2), 744–751. https://doi.org/10.1523/JNEUROSCI.13-02-00744.1993

Leutgeb, J. K., Leutgeb, S., Moser, M.-B., & Moser, E. I. (2007). Pattern Separation in the Dentate Gyrus and CA3 of the Hippocampus. Science, 315(5814), 961–966. https://doi.org/10.1126/science.1135801

Levy, S. J., Kinsky, N. R., Mau, W., Sullivan, D. W., & Hasselmo, M. E. (2021). Hippocampal spatial memory representations in mice are heterogeneously stable. Hippocampus, 31(3), 244– 260. https://doi.org/10.1002/hipo.23272

Liu, X., Ramirez, S., Pang, P. T., Puryear, C. B., Govindarajan, A., Deisseroth, K., & Tonegawa, S. (2012a). Optogenetic stimulation of a hippocampal engram activates fear memory recall. Nature, 484(7394), 381–385. https://doi.org/10.1038/nature11028

Marr, D. (1971). Simple memory: A theory for archicortex. Philosophical Transactions of the Royal Society of London. B, Biological Sciences, 262(841), 23–81. https://doi.org/10.1098/rstb.1971.0078

Mathis, A., Mamidanna, P., Cury, K. M., Abe, T., Murthy, V. N., Mathis, M. W., & Bethge, M. (2018). DeepLabCut: Markerless pose estimation of user-defined body parts with deep learning. Nature Neuroscience, 21(9), 1281–1289. https://doi.org/10.1038/s41593-018-0209-y

McLamb, R. L., Mundy, W. R., & Tilson, H. A. (1988). Intradentate colchicine disrupts the acquisition and performance of a working memory task in the radial arm maze. Neurotoxicology, 9(3), 521–528.

McNaughton, B. L., Barnes, C. A., Meltzer, J., & Sutherland, R. J. (1989). Hippocampal granule cells are necessary for normal spatial learning but not for spatially-selective pyramidal cell discharge. Experimental Brain Research, 76(3), 485–496. https://doi.org/10.1007/BF00248904

McNaughton, B. L., & Morris, R. G. M. (1987). Hippocampal synaptic enhancement and information storage within a distributed memory system. Trends in Neurosciences, 10(10), 408– 415. https://doi.org/10.1016/0166-2236(87)90011-7

Nakashiba, T., Cushman, J. D., Pelkey, K. A., Renaudineau, S., Buhl, D. L., McHugh, T. J., Barrera, V. R., Chittajallu, R., Iwamoto, K. S., McBain, C. J., Fanselow, M. S., & Tonegawa, S. (2012). Young Dentate Granule Cells Mediate Pattern Separation whereas Old Granule Cells Contribute to Pattern Completion. Cell, 149(1), 188–201. https://doi.org/10.1016/j.cell.2012.01.046

Nanry, K. P., Mundy, W. R., & Tilson, H. A. (1989). Colchicine-induced alterations of reference memory in rats: Role of spatial versus non-spatial task components. Behavioural Brain Research, 35(1), 45–53. https://doi.org/10.1016/S0166-4328(89)80007-5

Neunuebel, J. P., & Knierim, J. J. (2012). Spatial Firing Correlates of Physiologically Distinct Cell Types of the Rat Dentate Gyrus. Journal of Neuroscience, 32(11), 3848–3858. https://doi.org/10.1523/jneurosci.6038-11.2012

Neunuebel, J. P., & Knierim, J. J. (2014). CA3 Retrieves Coherent Representations from Degraded Input: Direct Evidence for CA3 Pattern Completion and Dentate Gyrus Pattern Separation. Neuron, 81(2), 416–427. https://doi.org/10.1016/j.neuron.2013.11.017

O’Keefe, J., & Dostrovsky, J. (1971). The hippocampus as a spatial map. Preliminary evidence from unit activity in the freely-moving rat. Brain Research, 34(1), 171–175. https://doi.org/10.1016/0006-8993(71)90358-1

O’Reilly, R. C., & McClelland, J. L. (1994). Hippocampal conjunctive encoding, storage, and recall: Avoiding a trade-off. Hippocampus, 4(6), 661–682. https://doi.org/10.1002/hipo.450040605

Pettit, N. L., Yap, E.-L., Greenberg, M. E., & Harvey, C. D. (2022). Fos ensembles encode and shape stable spatial maps in the hippocampus. Nature, 1–8. https://doi.org/10.1038/s41586-022-05113-1

Ramirez, S., Liu, X., Lin, P.-A., Suh, J., Pignatelli, M., Redondo, R. L., Ryan, T. J., & Tonegawa, S. (2013). Creating a False Memory in the Hippocampus. Science, 341(6144), 387–391. https://doi.org/10.1126/science.1239073

Redondo, R. L., Kim, J., Arons, A. L., Ramirez, S., Liu, X., & Tonegawa, S. (2014). Bidirectional switch of the valence associated with a hippocampal contextual memory engram. Nature, 513(7518), 426–430. https://doi.org/10.1038/nature13725

Reijmers, L. G., Perkins, B. L., Matsuo, N., & Mayford, M. (2007). Localization of a Stable Neural Correlate of Associative Memory. Science, 317(5842), 1230–1233. https://doi.org/10.1126/science.1143839

Roy, D. S., Arons, A., Mitchell, T. I., Pignatelli, M., Ryan, T. J., & Tonegawa, S. (2016). Memory retrieval by activating engram cells in mouse models of early Alzheimer’s disease. Nature, 531(7595), 508–512. https://doi.org/10.1038/nature17172

Santoro, A. (2013). Reassessing pattern separation in the dentate gyrus. Frontiers in Behavioral Neuroscience, 7. https://doi.org/10.3389/fnbeh.2013.00096

Sasaki, T., Piatti, V. C., Hwaun, E., Ahmadi, S., Lisman, J. E., Leutgeb, S., & Leutgeb, J. K. (2018). Dentate network activity is necessary for spatial working memory by supporting CA3 sharp-wave ripple generation and prospective firing of CA3 neurons. Nature Neuroscience, 21(2), 258–269. https://doi.org/10.1038/s41593-017-0061-5

Satvat, E., Schmidt, B., Argraves, M., Marrone, D. F., & Markus, E. J. (2011). Changes in Task Demands Alter the Pattern of zif268 Expression in the Dentate Gyrus. The Journal of Neuroscience, 31(19), 7163–7167. https://doi.org/10.1523/JNEUROSCI.0094-11.2011

Schmidt, U., Weigert, M., Broaddus, C., & Myers, G. (2018). Cell Detection with Star-Convex Polygons. In A. F. Frangi, J. A. Schnabel, C. Davatzikos, C. Alberola-López, & G. Fichtinger (Eds.), Medical Image Computing and Computer Assisted Intervention – MICCAI 2018 (pp. 265–273). Springer International Publishing. https://doi.org/10.1007/978-3-030-00934-2_30

Senzai, Y., & Buzsáki, G. (2017). Physiological properties and behavioral correlates of hippocampal granule cells and mossy cells. Neuron, 93(3), 691-704.e5. https://doi.org/10.1016/j.neuron.2016.12.011

Shen, J., Yao, P.-T., Ge, S., & Xiong, Q. (2021). Dentate granule cells encode auditory decisions after reinforcement learning in rats. Scientific Reports, 11(1), Article 1. https://doi.org/10.1038/s41598-021-93721-8

Straube, T., Korz, V., Balschun, D., & Uta Frey, J. (2003). Requirement of β-adrenergic receptor activation and protein synthesis for LTP-reinforcement by novelty in rat dentate gyrus. The Journal of Physiology, 552(Pt 3), 953–960. https://doi.org/10.1113/jphysiol.2003.049452

Straube, T., Korz, V., & Frey, J. U. (2003). Bidirectional modulation of long-term potentiation by novelty-exploration in rat dentate gyrus. Neuroscience Letters, 344(1), 5–8. https://doi.org/10.1016/S0304-3940(03)00349-5

Sun, X., Bernstein, M. J., Meng, M., Rao, S., Sørensen, A. T., Yao, L., Zhang, X., Anikeeva, P. O., & Lin, Y. (2020). Functionally Distinct Neuronal Ensembles within the Memory Engram. Cell, 181(2), 410-423.e17. https://doi.org/10.1016/j.cell.2020.02.055

Tanaka, K. Z., He, H., Tomar, A., Niisato, K., Huang, A. J. Y., & McHugh, T. J. (2018). The hippocampal engram maps experience but not place. Science. https://doi.org/10.1126/science.aat5397

Treves, A., & Rolls, E. T. (1994). Computational analysis of the role of the hippocampus in memory. Hippocampus, 4(3), 374–391. https://doi.org/10.1002/hipo.450040319

VanElzakker, M., Fevurly, R. D., Breindel, T., & Spencer, R. L. (2008). Environmental novelty is associated with a selective increase in Fos expression in the output elements of the hippocampal formation and the perirhinal cortex. Learning & Memory, 15(12), 899–908. https://doi.org/10.1101/lm.1196508

Weigert, M., Schmidt, U., Haase, R., Sugawara, K., & Myers, G. (2020). Star-convex Polyhedra for 3D Object Detection and Segmentation in Microscopy. 2020 IEEE Winter Conference on Applications of Computer Vision (WACV), 3655–3662. https://doi.org/10.1109/WACV45572.2020.9093435

Wood, E. R., Dudchenko, P. A., Robitsek, R. J., & Eichenbaum, H. (2000). Hippocampal Neurons Encode Information about Different Types of Memory Episodes Occurring in the Same Location. Neuron, 27(3), 623–633. https://doi.org/10.1016/S0896-6273(00)00071-4

Xavier, G. F., & Costa, V. C. I. (2009). Dentate gyrus and spatial behaviour. Progress in Neuro-Psychopharmacology and Biological Psychiatry, 33(5), 762–773. https://doi.org/10.1016/j.pnpbp.2009.03.036

Xavier, G. F., Oliveira-Filho, F. J. B., & Santos, A. M. G. (1999). Dentate gyrus-selective colchicine lesion and disruption of performance in spatial tasks: Difficulties in “place strategy” because of a lack of flexibility in the use of environmental cues? Hippocampus, 9(6), 668–681. https://doi.org/10.1002/(SICI)1098-1063(1999)9:6<668::AID-HIPO8>3.0.CO;2-9

Zaki, Y., Mau, W., Cincotta, C., Monasterio, A., Odom, E., Doucette, E., Grella, S. L., Merfeld, E., Shpokayte, M., & Ramirez, S. (2022). Hippocampus and amygdala fear memory engrams re-emerge after contextual fear relapse. Neuropsychopharmacology, 47(11), 1992–2001. https://doi.org/10.1038/s41386-022-01407-0

